# Tau-Associated Neuronal Loss in the Intermediate Nucleus of the Human Hypothalamus (VLPO Analog): Unveiling the Basis of NREM Sleep Dysfunction in PSP and Alzheimer’s Disease

**DOI:** 10.1101/2025.03.11.642707

**Authors:** Shima Rastegar-Pouyani, Caroline Lew, Felipe L. Pereira, Abhijit Satpati, Vitor Paes, Renata Elaine Paraízo Leite, Claudia Kimie Suemoto, Salvatore Spina, Helmut Heinsen, William W. Seely, Christine M. Walsh, Thomas C. Neylan, Lea T. Grinberg

## Abstract

Sleep disturbances are prevalent in Alzheimer’s disease (AD) and Progressive Supranuclear Palsy (PSP), often exacerbating disease progression. Understanding the neuropathological basis of these disturbances is essential for identifying potential therapeutic targets. This study investigates the intermediate nucleus (IntN) of the human hypothalamus—a key sleep-regulating region analogous to the rodent ventrolateral preoptic area (VLPO)—to assess neuronal loss and tau pathology in AD and PSP. Using postmortem brain tissue, we applied unbiased stereology to quantify galanin-expressing neurons and phosphorylated tau (p-tau) accumulation.

Among 26 cases analyzed, both AD and PSP exhibited significant neuronal loss in the IntN, with PSP showing the most pronounced reduction (84.9% fewer neurons than healthy controls [HC]). In AD, neuronal loss correlated with Braak staging, with late-stage AD cases (Braak 5–6) demonstrating a 76.9% reduction in galanin-expressing neurons compared to HC, while non-galanin neurons exhibited a more moderate decline (45.7%). In PSP, extensive neuronal loss precluded a clear assessment of p-tau burden. These findings suggest a differential neuronal vulnerability to tau pathology across diseases, aligning with distinct sleep disturbances observed in each condition. PSP, characterized by severe insomnia despite preserved wake-promoting neurons, may be explained by the near-total loss of NREM sleep-regulating neurons. In contrast, AD exhibits a progressive decline in both wake- and sleep-promoting neurons, contributing to excessive daytime sleepiness and sleep fragmentation.

This study provides critical insights into the selective neuronal vulnerabilities underlying sleep dysfunction in tauopathies, emphasizing the need for targeted interventions to mitigate sleep disturbances in these disorders.

## Introduction

Sleep disturbances are common in neurodegenerative diseases and often present distinct, disease-specific profiles (Iranzo, 2016). Among tauopathies, conditions marked by the pathological accumulation of tau protein in neuronal populations, progressive supranuclear palsy (PSP), and Alzheimer’s disease (AD) exhibit particularly divergent sleep phenotypes. In PSP, patients experience severe insomnia-like symptoms, characterized by persistent difficulty initiating sleep without compensatory daytime sleepiness, along with hyperarousal and reduced sleep drive (Höglinger et al., 2017)(Walsh et al., 2017). In contrast, AD, the most prevalent tauopathy, presents subtype-specific sleep disturbances. In typical (amnestic) AD cases, memory impairment is the initial and dominant feature, and these patients often show significant slow-wave sleep (SWS) loss with relatively preserved REM sleep (Moran et al., 2005)(Ju et al., 2014)(Polsinelli & Apostolova, 2022). Atypical AD cases, however, can manifest with non-memory deficits early on and more intact SWS patterns, although REM sleep may be reduced (Falgàs et al., 2023; Walsh et al., 2017)(Pina Escudero et al., 2024; Wang et al., 2024).

Despite growing recognition of these distinct sleep signatures, the underlying neurobiological mechanisms remain poorly understood. The regulation of sleep and wakefulness involves a complex interplay of interconnected neuronal populations in the hypothalamus and brainstem. These include wake-promoting nuclei, REM- and NREM-promoting nuclei (Saper & Fuller, 2017). Given that PSP patients have difficulty initiating and maintaining sleep, we hypothesized that the profound inability to generate SWS in PSP may arise from a dramatic loss of sleep-promoting neurons with relative preservation of wake-promoting neurons in contrast to AD, where both wake, and sleep-regulating circuits are compromised. Evidence suggests that AD is associated with severe degeneration of wake-promoting neurons, while these populations remain relatively preserved in PSP (Horner & Peever, 2017)(Arendt et al., 2015)(Kaalund et al., 2020)(J. Oh, Eser, et al., 2019)(Satpati et al., 2023). Conversely, changes in sleep-promoting nuclei remain largely unexplored in tauopathies.

A critical node in the sleep-wake regulatory network is the intermediate nucleus (IntN) of the human hypothalamus, considered analogous to the ventrolateral preoptic area (VLPO) in rodents (Saper, 2021; Tsuneoka & Funato, 2021). The IntN is characterized by a cluster of medium-sized, round neurons mostly identifiable by galanin immunoreactivity (Gai et al., 1990)(Swaab & Hofman, 1988). Galanin-positive neurons in the VLPO have been shown in animal models to promote and maintain NREM sleep—especially SWS—by providing inhibitory input to wake-active neuronal populations (Kroeger et al., 2018)(Szymusiak et al., 1998).Studies in humans indicate that reductions in IntN galanin neurons correlate with sleep fragmentation and diminished sleep quality in advanced age and AD (Lim et al., 2014; Saper, 2021). However, most prior human data have focused on aging, sexual dimorphisms, or normal variation (Garcia Falgueras et al., 2011) (Swaab & Fliers, 1985), rather than disease-specific vulnerabilities across distinct tauopathies.

Galanin-positive neurons in the ventrolateral preoptic nucleus (VLPO) promote and maintain NREM sleep, especially slow-wave sleep (SWS), by inhibiting wake-active neurons (Kroeger et al., 2018; Szymusiak et al., 1998). About 70% of galanin neurons in the preoptic area are GABAergic, suggesting they can inhibit several brainstem and hypothalamic nuclei, including the laterodorsal tegmental nucleus (LDT), dorsal raphe nucleus (DRN), median raphe nucleus (MRN), locus coeruleus (LC), and ventrolateral periaqueductal gray (vlPAG) (Steininger et al., 2001; Lu et al., 2002; (J. Oh, Petersen, et al., 2019). VLPO neurons also receive input from regions such as the tuberomammillary nucleus (TMN), raphe nuclei, LC, and hypothalamic areas like the median preoptic area (MnPO) and lateral hypothalamus (LH) (Chou et al., 2002).

In this study, we leveraged well-characterized postmortem human brains from individuals with PSP, amnestic AD, atypical AD, and age-matched healthy controls to quantify total neuronal populations in the IntN, and galaninergic- and tau-positive neuronal subpopulations. Based on the pattern of sleep dysfunction, we hypothesize that IntN degeneration will be worse in PSP, followed by amnestic AD and atypical AD variants. By determining whether the IntN’s sleep-promoting neurons are disproportionately lost or affected by tau pathology in PSP and specific AD variants, we aimed to establish a mechanistic framework that clarifies how selective neuronal vulnerabilities contribute to disease-specific sleep phenotypes. This work sought to link the degeneration of hypothalamic sleep-regulatory centers to the distinct sleep disturbances observed in these tauopathies. Ultimately, this study has the potential to provide a foundation for developing effective interventions targeting sleep dysfunction in neurodegenerative diseases.

## Materials and Methods

This study was deemed non-human by the Institutional Review Board. We analyzed postmortem human hypothalami sourced from two cohorts: the Neurodegenerative Disease Brain Bank (NDBB) at the University of California, San Francisco, and the Brazilian Biobank for Aging Studies (BBAS) at the University of São Paulo (Grinberg et al., 2007)(Spina et al., 2021; Suemoto et al., 2017). Neuropathological evaluations followed the established international protocols and diagnostic criteria.

Inclusion criteria required participants to be over 50 years of age, with no history of neurological or primary psychiatric disorders unrelated to the main diagnosis. The following diagnostic categories were included: (1) Alzheimer’s Disease (AD) neuropathological changes: Any level of pathology, provided no other primary or contributing neuropathological diagnosis was present. AD cases were categorized as follows: a) Mid-stage AD: Braak stage 3 or 4 (Braak & Braak, 1991), Thal phase 1 or greater (Thal et al., 2002); b) Late-stage AD: Braak stage 5 or 6, Thal phase 1 or greater. Clinical data from skilled clinicians were used to classify AD cases as typical (amnestic) or atypical variants. No mid-stage AD cases met the criteria for atypical presentations (2) Progressive Supranuclear Palsy (PSP): Confirmed PSP pathology with Braak stage ≤ 4 for AD pathology and absence of other primary neurodegenerative diagnoses. (3) Control cases: Defined as Braak stage 0-2, Thal 0-1, no non-AD neuropathology diagnosis and no cognitive decline.

The presence of Lewy body pathology was an exclusion criterion, except for cases where Lewy bodies were confined to the amygdala (amygdala-predominant)(Attems et al., 2021), as these cases lack Lewy bodies in the hypothalamic axis (data not shown). Any TDP-43 proteinopathy was also an exclusion criterion.

Functional cognitive measures were available for all cases, assessed through the Clinical Dementia Rating (CDR) scale, reflecting cognitive status within three months prior to patient death(Ehrenberg et al., 2023).

### Project -Specific Tissue Processing and Immunohistochemistry

Previous studies have highlighted the difficulty in consistently delineating the IntN in Nissl-stained human hypothalami (Brockhaus, 1942)(Swaab & Hofman, 1988). To address this, we combined cytoarchitectonic and immunohistochemical approaches. Formalin-fixed hypothalamic blocks encompassing the entire IntN were embedded in celloidin, serially sectioned at 30 μm thickness, and collected in coronal or horizontal planes (Heinsen et al., 2000; Theofilas et al., 2014).

The IntN, located near the crossing of the anterior commissure and lateral to the supraoptic (SON) and suprachiasmatic nuclei, was identified using both gross anatomical landmarks and histological cues. Following established definitions (Allen et al., 1989)(Swaab & Hofman, 1988)(Lim et al., 2014), we located the IntN as a cluster of medium-sized, round neurons (**Figure 1**). Every second section through the region of interest (ROI) was double-immunostained overnight at room temperature for galanin (MAB5854, R&D Systems, 1:300) and phosphorylated tau (p-tau T231, ab151559, Abcam, 1:3000), and then counterstained with gallocyanin (pH 2.1) to visualize all neurons. Galanin immunoreactivity served as a reliable chemical marker to define IntN borders more objectively, overcoming previous limitations related to subjective boundary delineation (Gai et al., 1990) (Garcia Falgueras et al., 2011). Cytoarchitectonic landmarks, including magnocellular cells in the supraoptic area and galanin-positive neurons, were used to histologically locate the region of interest.

**Figure 1.**
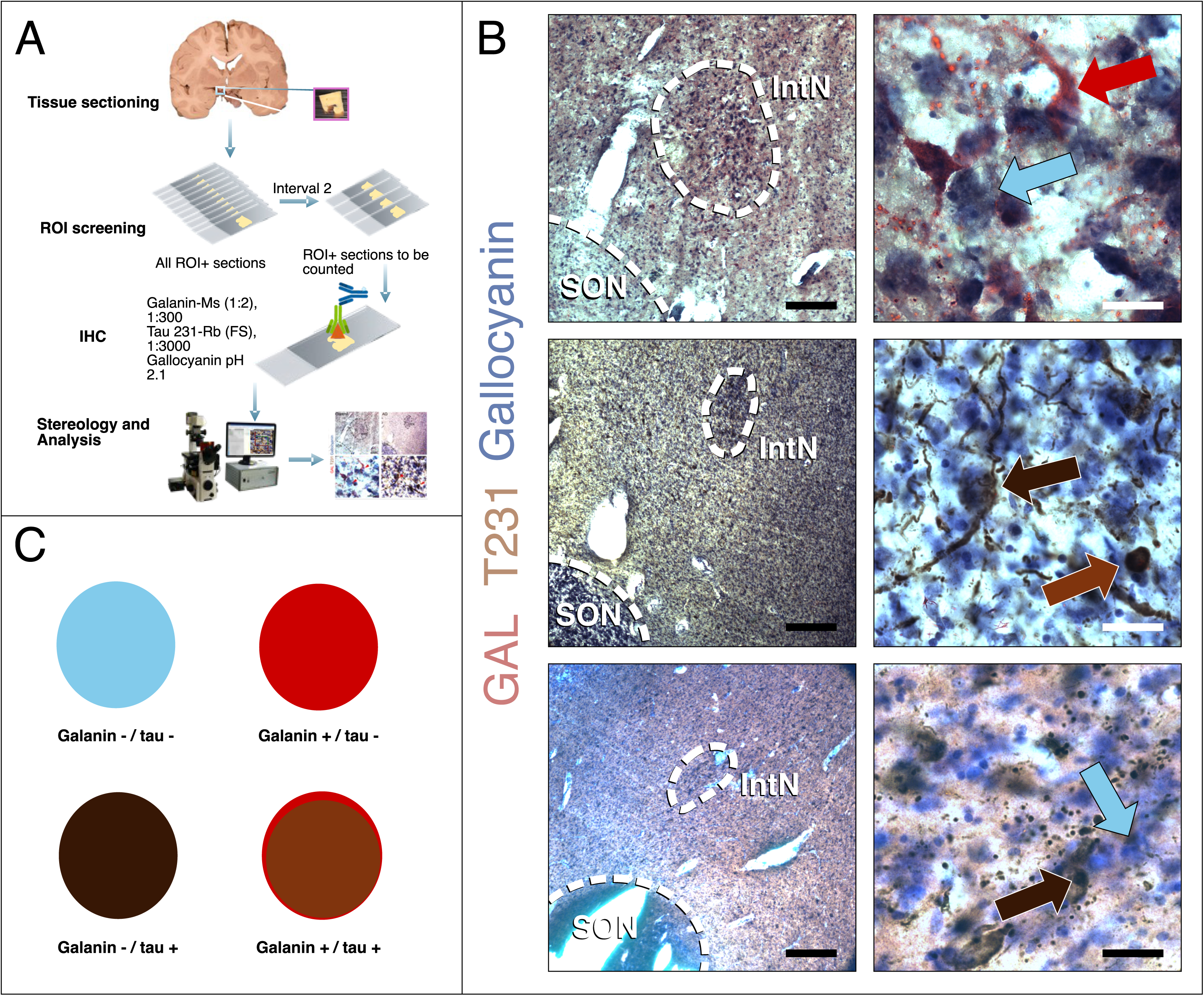
Neuronal counts and neuronal tau burden in galanin-positive and galanin-negative neurons of the IntN in AD and PSP. (A) Summary of the tissue processing and stereology workflow. (B) Representative microphotographs of the histological section containing the preoptic area of the hypothalamus of postmortem human tissue in AD, PSP, and healthy controls. The IntN neurons are distinguishable from the rest of the preoptic region by their medium size and their arrangement as a compact, rounded cluster of neurons, indicated by the traced lines in the upper row. The right column micrographs show examples of neurons expressing galanin (GAL) or not expressing galanin and containing or not containing p-tau inclusions (T231). Blue arrow: Galanin - / tau - neuron; Dark brown arrow: Galanin - / tau + neuron; Red arrow: Galanin + / tau - neuron; Reddish-brown (or burnt orange) arrow: Galanin + / tau + neuron. (C) Scheme of the neuronal subgrouping for statistical analysis.

For immunohistochemical staining, free-floating hypothalamic sections were treated with 0.3% hydrogen peroxide in methanol to quench endogenous peroxidase activity, followed by a 50- minute incubation at 97°C in 0.01 M citrate buffer containing 0.1% Triton X-100 in 1X PBS (pH 6.0) for epitope retrieval. To minimize non-specific staining, sections were blocked for 40 minutes in a buffer containing 5% milk in PBST (0.1% Triton X). Serial sections were double-stained overnight at room temperature with monoclonal anti-galanin antibody (1:300, mouse, MAB5854, Biotechne R&D System, MN, USA) and monoclonal anti-Tau phosphorylated at Threonine 231 (1:3000, rabbit, ab151559, Abcam, Cambridge, UK). After primary antibody incubation, sections were incubated with HRP-conjugated anti-rabbit IgG (1:400, Advansta, CA, USA) for 1 hour at room temperature. T231 staining was visualized using the ImmPACT DAB Peroxidase Substrate Kit (Vector Labs, CA, USA), while Galanin was visualized using the MACH 2 Mouse AP Polymer (Biocare Medical, CA, USA) and Vector Red Substrate Kit (Vector Labs, CA, USA). Sections were counterstained with gallocyanin (Alfa Aesar, MA, USA) at an optimized pH of 2.1 to stain nucleic acids and visualize all neurons for neuronal quantification. (Garcia Falgueras et al., 2011)

### Unbiased Stereology for Estimating Neuronal Populations

To accurately estimate neuronal populations, we employed the optical fractionator probe in Stereo Investigator v.10 (MBF Bioscience, VT, USA) with live images captured via a high-resolution camera attached to an Axio Imager.A2 microscope (Carl Zeiss Microscopy, NY, USA) - a method not influenced by cutting plane and well-established for human neuropathological assessments (Theofilas et al., 2014).

The ROI was traced under low magnification (5x), and neuronal counts were performed at 63x (oil immersion). Stereological parameters were determined using the “resample–oversample” method (Slomianka & West, 2005; West, 2013), with a 6 μm guard zone and a 14 μm dissector height (details in Supplementary Table S1). We calculated coefficients of error (CE) to ensure sampling precision and values under 0.1 were considered acceptable. Each neuron was classified by galanin expression and tau pathology (Figure 1). The volume of the IntN was calculated using the Cavalieri method using the Stereoinvestigator setup (Theofilas et al., 2018).

### Statistical Analyses

Demographic differences were analyzed using chi-square, one-way ANOVA, Welch’s ANOVA, or the Kruskal-Wallis tests, based on the data type, normality, and homogeneity of variance. Variations in neuronal counts and the proportions of neuronal subcategories were evaluated using the pairwise Wilcoxon rank-sum test, with p-values adjusted for multiple comparisons using the Benjamini-Hochberg method. Additionally, a mixed linear regression model was employed to account for the effects of sex, age at death (AaD), and post-mortem interval (PMI) on the comparisons (*e.g.*, *m*0 = measure ∼ group + Sex + AaD + PMI + (1 | Subject)). Considering the rare nature of the subjects, all available cases were used in the analysis’s comparison. A posterior power analysis simulation (1000× bootstrap) was conducted using each measure mean and standard deviation as key to generating uniform random values. The alpha level was set at 5%, and statistical power was defined as a simulated power analysis at 80%. The functions *R::cor* and *R::ggcorrplot::cor_pmat* were utilized to calculate Spearman correlation coefficients and their corresponding p-values. Z-score values were computed by subtracting each value from the mean of the HC group and dividing the result by the standard deviation of the HC group. To compare data exclusively from mid- and late-AD cases as a function of their Braak stage, a mixed linear regression model was applied. This model accounted for the same confounding variables as *m*0, with the case group variable replaced by Braak stage (*e.g.*, *m*1 = measure ∼ Braak Stage + Sex + AaD + PMI + (1 | Subject)). All analyses were conducted using R software (version 4.2.2; R Foundation for Statistical Computing, Vienna, Austria).

## Results

Of the 65 cases screened, 30 were excluded due to partial or complete loss of the ROI, nine for low tissue quality, and two for not meeting all neuropathological criteria. This resulted in a final sample of 26 cases. The average age at death of participants was 68.9 ± 10.0 years, with 61.5% being male. There were no significant differences in age at death, sex, disease duration, brain weight, or PMI distribution between the groups. Participant demographics are summarized in Table 1, and detailed neuropathological and demographic information for each participant is listed in Tables S1 and S2. Table 2 displays the absolute and relative counts of the neuronal (sub)population.

**Table 1.** Demographic, clinical, and neuropathological characteristics stratified by disease group.

| Characteristics | HC | mid-AD | late-AD | PSP | p-value |
| --- | --- | --- | --- | --- | --- |
| N | 6 | 4 | 8 | 8 |  |
| Age at Death, mean (SD), y | 66.8 (10.68) | 72.2 (6.4) | 69.1 (13.6) | 68.6 (7.91) | 0.883<sup>A</sup> |
| Males, No. (%) | 4 (66.7) | 2 (50) | 4 (50) | 6 (75) | 0.716<sup>C</sup> |
| Disease duration, mean (SD), y | 0 (0) | 7 (NA) | 8.5 (3.62) | 8.8 (3.41) | 0.844<sup>K</sup> |
| Braak Stage, range | 1-2 | 3-4 | 5-6 | 0-3 | <0.001<sup>C</sup> |
| CDR-SOB, median (IQR) | 0 (0) | 0 (4) | 17 (4.5) | 12 (4.5) | 0.003<sup>K</sup> |
| Brain weight, mean (SD), g | 1123.7 (580.75) | 1263.2 (109.06) | 1118.6 (141.18) | 1222.2 (203.92) | 0.403<sup>K</sup> |
| PMI, mean (SD), h | 14.9 (4.33) | 18.4 (6.9) | 14 (6.77) | 11.5 (7.53) | 0.442<sup>A</sup> |

Abbreviations: HC – Healthy controls; mid-AD – Alzheimer’s disease mid-stage cases; late-AD – Alzheimer’s disease late-stage cases; PSP – Progressive supranuclear palsy; N – number of individuals; SD – standard deviation; IQR – interquartile range; PMI – postmortem interval; NA – not accessed. P-values were calculated conforming data type, normality, and homogeneity of variance: A – One-way ANOVA; C – chi-square; K – Kruskal-Wallis.

**Table 2.**
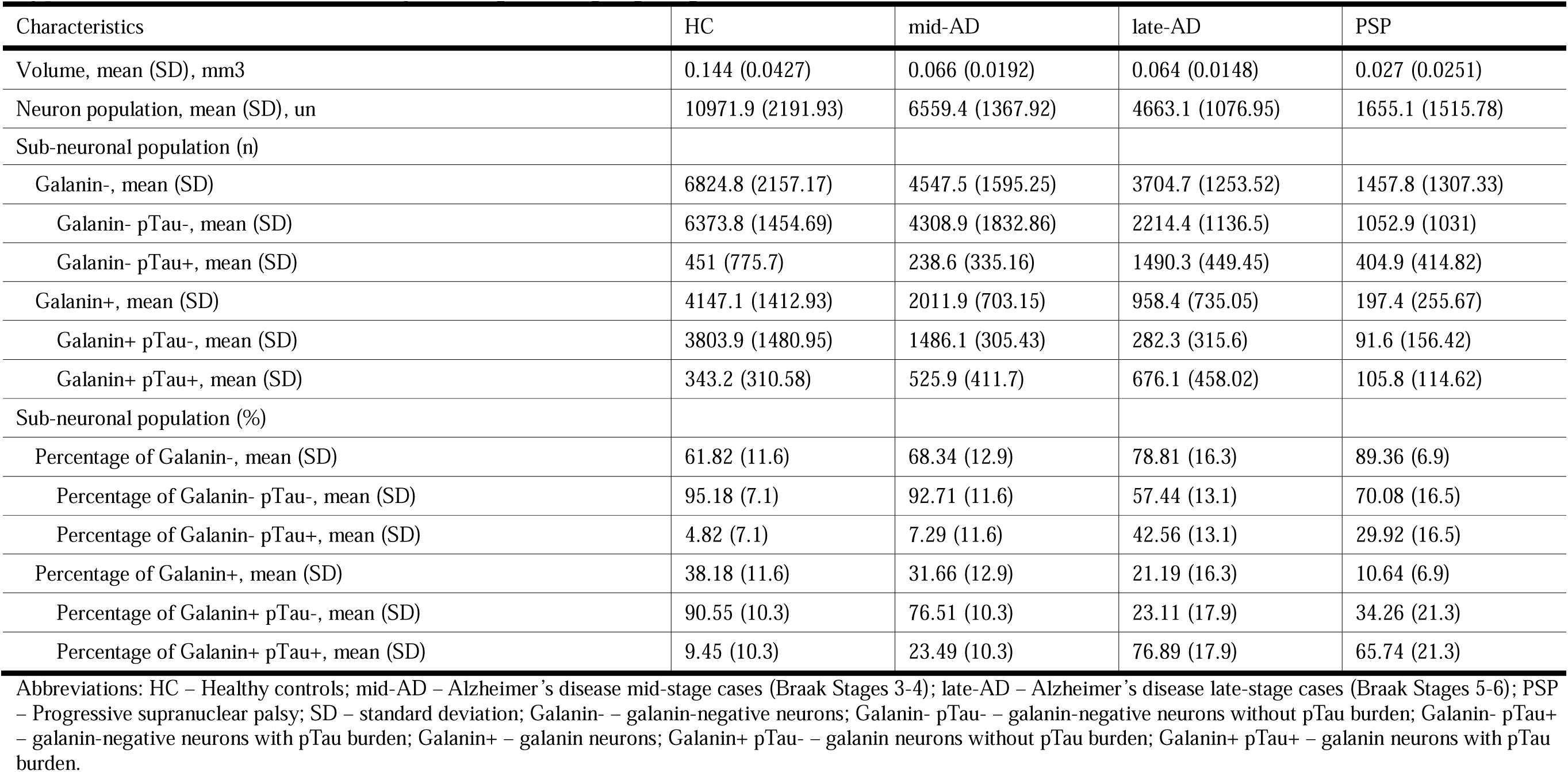
Mean (SD) and percentage of stereological estimates for different characteristics of the *intermediate nucleus of hypothalamus* (IntN) stratified by neuropathologic group.

| Characteristics | HC | mid-AD | late-AD | PSP |
| --- | --- | --- | --- | --- |
| Volume, mean (SD), mm3 | 0.144 (0.0427) | 0.066 (0.0192) | 0.064 (0.0148) | 0.027 (0.0251) |
| Neuron population, mean (SD), un | 10971.9 (2191.93) | 6559.4 (1367.92) | 4663.1 (1076.95) | 1655.1 (1515.78) |
| Sub-neuronal population (n) |  |  |  |  |
| Galanin-, mean (SD) | 6824.8 (2157.17) | 4547.5 (1595.25) | 3704.7 (1253.52) | 1457.8 (1307.33) |
| Galanin- pTau-, mean (SD) | 6373.8 (1454.69) | 4308.9 (1832.86) | 2214.4 (1136.5) | 1052.9 (1031) |
| Galanin- pTau+, mean (SD) | 451 (775.7) | 238.6 (335.16) | 1490.3 (449.45) | 404.9 (414.82) |
| Galanin+, mean (SD) | 4147.1 (1412.93) | 2011.9 (703.15) | 958.4 (735.05) | 197.4 (255.67) |
| Galanin+ pTau-, mean (SD) | 3803.9 (1480.95) | 1486.1 (305.43) | 282.3 (315.6) | 91.6 (156.42) |
| Galanin+ pTau+, mean (SD) | 343.2 (310.58) | 525.9 (411.7) | 676.1 (458.02) | 105.8 (114.62) |
| Sub-neuronal population (%) |  |  |  |  |
| Percentage of Galanin-, mean (SD) | 61.82 (11.6) | 68.34 (12.9) | 78.81 (16.3) | 89.36 (6.9) |
| Percentage of Galanin- pTau-, mean (SD) | 95.18 (7.1) | 92.71 (11.6) | 57.44 (13.1) | 70.08 (16.5) |
| Percentage of Galanin- pTau+, mean (SD) | 4.82 (7.1) | 7.29 (11.6) | 42.56 (13.1) | 29.92 (16.5) |
| Percentage of Galanin+, mean (SD) | 38.18 (11.6) | 31.66 (12.9) | 21.19 (16.3) | 10.64 (6.9) |
| Percentage of Galanin+ pTau-, mean (SD) | 90.55 (10.3) | 76.51 (10.3) | 23.11 (17.9) | 34.26 (21.3) |
| Percentage of Galanin+ pTau+, mean (SD) | 9.45 (10.3) | 23.49 (10.3) | 76.89 (17.9) | 65.74 (21.3) |

Abbreviations: HC – Healthy controls; mid-AD – Alzheimer’s disease mid-stage cases (Braak Stages 3-4); late-AD – Alzheimer’s disease late-stage cases (Braak Stages 5-6); PSP – Progressive supranuclear palsy; SD – standard deviation; Galanin- – galanin-negative neurons; Galanin- pTau- – galanin-negative neurons without pTau burden; Galanin- pTau+ – galanin-negative neurons with pTau burden; Galanin+ – galanin neurons; Galanin+ pTau- – galanin neurons without pTau burden; Galanin+ pTau+ – galanin neurons with pTau burden.

To ensure quality control and assess the alignment of age at death between disease groups and HC, we initially generated a Spearman correlation plot as part of the exploratory analysis (Figure 2). Across all cases, and within the PSP group, PMI and age at death showed no correlation with other phenotypic variables or neuronal counts. We found a negative correlation between the Braak stages and the total number of neurons in AD cases (ρ = −0.7, p = 0.009); where, notably, the count of galanin-negative neurons showed no significant correlation, whereas galanin-positive neurons exhibited a strong negative correlation (ρ = −0.6, p = 0.040). Additionally, expected disease duration and CDR-SOB correlations were detected between each other, CDR-SOB versus Braak, as well as with neuronal counts, as illustrated in Figure 2.

**Figure 2.**
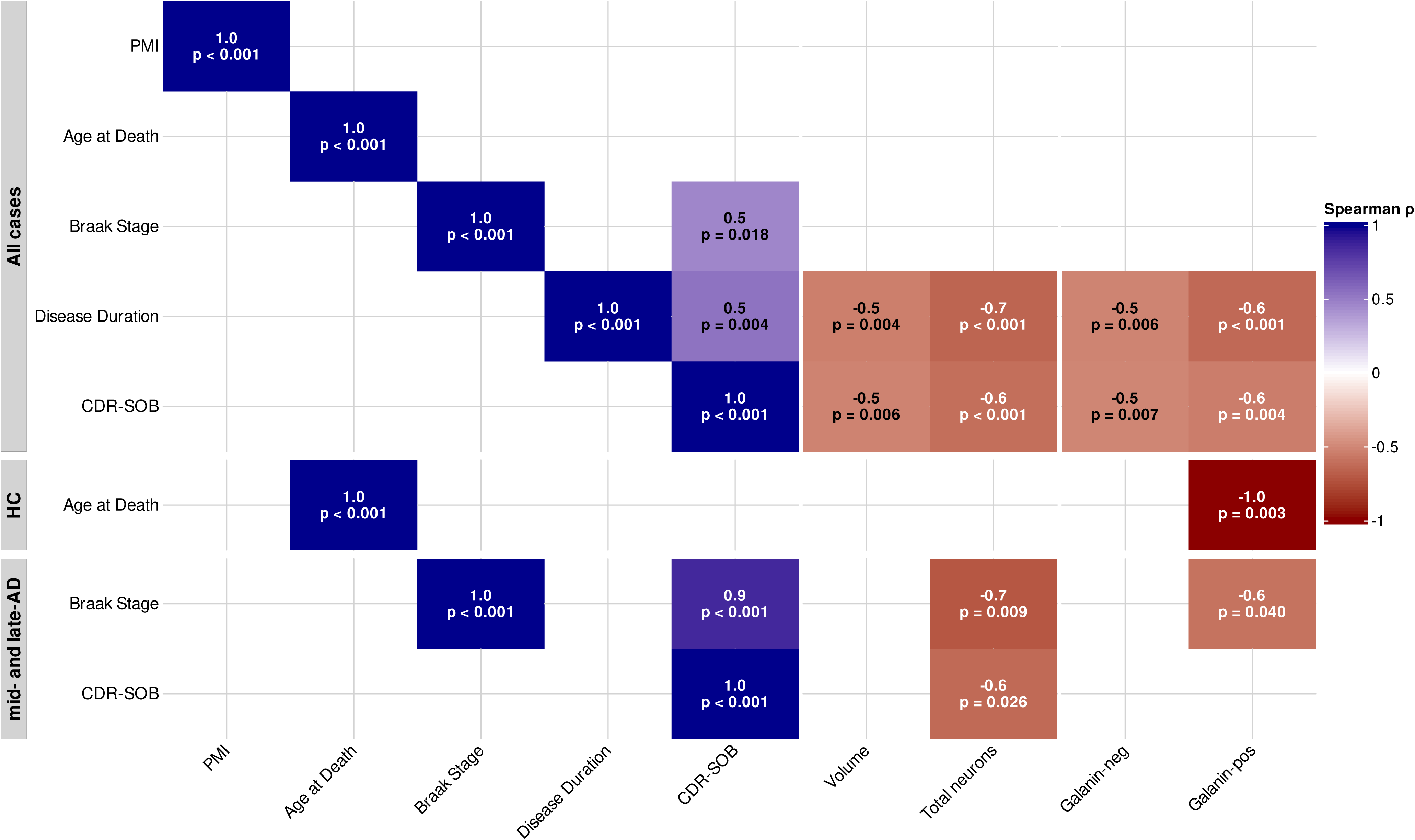
Correlation matrix between phenotypic and neuropathological measurements and postmortem stereological neuronal counts. Darker blue shades represent stronger positive correlations, while darker red shades indicate stronger negative correlations. Only statistically significant correlations (*p* < 0.05) are displayed as colored blocks. The “All Cases” group includes 26 samples; the HC group consists of 6 samples and the mid- and late-AD groups include 12 samples. No significant correlations were detected in the PSP group, which is therefore excluded from the figure.

Next, we applied a non-parametric test and mixed linear model to compare the stereologically determined neuronal estimates for each group. Table 3 presents the adjusted analyses for each group comparison, along with the *a posteriori* power analyses. The comparison’s significance and power were similar in both analytical methods.

**Table 3.** Associations between volume and different stereological estimates with neuropathologic groups.

|  | HC vs. mid-AD | HC vs. late-AD | HC vs. PSP | mid-AD vs. late-AD | Amn-late-AD vs. Atyp-late-AD | PSP vs. mid-AD | PSP vs. late-AD |
| --- | --- | --- | --- | --- | --- | --- | --- |
| Wilcoxon rank<sup>1</sup> test, p-adjusted<sup>2</sup>; power |  |  |  |  |  |  |  |
| Volume | 0.0571; 57.4 | 0.004; 77.6 | **0.0115; 95.3** | 0.933; 3.4 | 1; 4.6 | 0.065; 39.1 | 0.0394; 69.7 |
| Neuron population | 0.0241; 70.3 | **0.004; 99.2** | **0.00944; 100** | 0.0485; 48.9 | 1; 3.1 | **0.0241; 95.8** | **0.00944; 93.9** |
| Sub-neuronal population (n) |  |  |  |  |  |  |  |
| Galanin- | 0.133; 14.2 | 0.0233; 52.1 | **0.0139; 93.6** | 0.683; 6.3 | 0.393; 3.5 | 0.0394; 54.8 | 0.0394; 62.1 |
| Galanin- pTau- | 0.145; 14.8 | **0.004; 95.9** | **0.0115; 99.7** | 0.145; 30.9 | 0.786; 4 | 0.0838; 57.5 | 0.145; 25.8 |
| Galanin- pTau+ | 1; 1.2 | 0.117; 37.9 | 1; 2.3 | **0.0202; 86.2** | 0.25; 3.5 | 1; 1.7 | **0.0113; 90.5** |
| Galanin+ | 0.0381; 40.3 | **0.004; 89.4** | **0.0115; 97.5** | 0.0727; 32.1 | 0.393; 3.9 | **0.0289; 95.8** | 0.0289; 50 |
| Galanin+ pTau- | 0.0241; 60.8 | **0.004; 94.9** | **0.0115; 96.1** | **0.0162; 98.8** | 0.571; 3.7 | **0.0241; 100** | 0.139; 22.8 |
| Galanin+ pTau+ | 0.705; 3.6 | 0.42; 7.8 | 0.42; 16.7 | 0.705; 1.8 | 0.393; 4 | 0.105; 34.4 | 0.0161; 48.7 |
| Sub-neuronal population (%) |  |  |  |  |  |  |  |
| Percentage of Galanin- | 0.848; 2.3 | 0.325; 19.9 | **0.026; 90.6** | 0.848; 4 | 0.571; 4.6 | 0.159; 51.4 | 0.848; 13.3 |
| Percentage of Galanin- pTau- | 1; 2.8 | 0.466; 15.9 | 1; 2.2 | 0.436; 13.3 | 0.786; 3.5 | 1; 3.6 | 0.466; 27.1 |
| Percentage of Galanin- pTau+ | 1; 1.7 | **0.004; 96.1** | 0.0346; 57.3 | **0.0202; 88.5** | 0.571; 3.6 | 0.0952; 45.5 | 1; 4.2 |
| Percentage of Galanin+ | 0.848; 3.4 | 0.325; 20.3 | **0.026; 90.7** | 0.848; 4.9 | 0.571; 3.8 | 0.159; 50.6 | 0.848; 13.6 |
| Percentage of Galanin+ pTau- | 0.343; 15.2 | **0.00799; 89.3** | **0.0216; 93.2** | 0.0323; 75.7 | 0.571; 3 | **0.0476; 88.5** | 0.943; 2.8 |
| Percentage of Galanin+ pTau+ | 0.19; 13.5 | 0.016; 37.9 | 0.208; 36.3 | 0.566; 4.7 | 0.25; 2.5 | 1; 8.2 | 0.28; 27.1 |
| Mixed linear model<sup>3</sup>, p-value; power |  |  |  |  |  |  |  |
| Volume | 0.0121; 67.7 | **0.00394; 94.3** | **3.26e-05; 99.2** | 0.855; 6.1 | 0.37; 5.5 | 0.0426; 43.5 | 0.0013; 74 |
| Neuron population | 0.0439; 66.1 | **0.000278; 100** | **6.67e-05; 100** | 0.0729; 49 | 0.674; 4.8 | **0.00466; 94.1** | **0.000541; 90.6** |
| Sub-neuronal population (n) |  |  |  |  |  |  |  |
| Galanin- | 0.29; 24 | 0.0216; 73.1 | **0.000834; 98.3** | 0.455; 13.3 | 0.696; 5.7 | 0.0195; 60.9 | 0.00826; 71.7 |
| Galanin- pTau- | 0.295; 23.5 | **0.000531; 99.6** | **0.000147; 100** | 0.068; 44.6 | 0.953; 5.7 | 0.0185; 68.1 | 0.0365; 34.5 |
| Galanin- pTau+ | 0.439; 5.8 | 0.0177; 65.4 | 0.977; 6 | **0.00277; 93.8** | 0.249; 6.6 | 0.668; 8.8 | **0.00156; 93.9** |
| Galanin+ | 0.0247; 44.3 | **0.00166; 97.3** | **0.000228; 99.9** | 0.192; 45.7 | 0.257; 5.4 | **0.00544; 95.2** | 0.0559; 51.2 |
| Galanin+ pTau- | 0.0233; 57.3 | **0.000463; 99.4** | **0.00039; 99.2** | **0.00323; 99.2** | 0.388; 6.1 | **0.000768; 100** | 0.219; 19.1 |
| Galanin+ pTau+ | 0.854; 10.5 | 0.129; 24.3 | 0.147; 29.6 | 0.434; 6.9 | 0.196; 4.6 | 0.128; 42.8 | 0.0256; 70.6 |
| Sub-neuronal population (%) |  |  |  |  |  |  |  |
| Percentage of Galanin- | 0.439; 9.3 | 0.165; 39.7 | **0.045; 97.2** | 0.625; 13 | 0.314; 4.2 | 0.98; 62.9 | 0.46; 24.8 |
| Percentage of Galanin- pTau- | 0.305; 11.9 | 0.023; 32.6 | 0.821; 7.7 | 0.0608; 29.2 | 0.773; 5.5 | 0.182; 8.4 | 0.131; 44.5 |
| Percentage of Galanin- pTau+ | 0.544; 5.7 | **0.000955; 99.4** | 0.0834; 74.8 | **0.00287; 93.4** | 0.00636; 4.4 | 0.151; 49.3 | 0.549; 9.3 |
| Percentage of Galanin+ | 0.439; 8.3 | 0.165; 41.5 | **0.045; 96.7** | 0.625; 12.5 | 0.314; 5.2 | 0.98; 61.4 | 0.46; 24.6 |
| Percentage of Galanin+ pTau- | 0.251; 20.8 | **0.00285; 97.6** | **0.0311; 97.8** | **0.0275; 84.4** | 0.465; 6 | **0.285; 92.4** | 0.926; 7.2 |
| Percentage of Galanin+ pTau+ | 0.291; 28.4 | 0.0337; 70 | 0.103; 52.8 | 0.265; 15.7 | 0.238; 4.6 | 0.282; 14.1 | 0.264; 48.1 |

Abbreviations: HC – Healthy controls; mid-AD – Alzheimer’s disease mid-stage cases (Braak Stages 3-4); late-AD – Alzheimer’s disease late-stage cases (Braak Stages 5-6); Amn-late-AD – Amnestic Alzheimer’s disease late-stage cases (Braak Stages 5-6); Atyp-late-AD – Atypical Alzheimer’s disease late-stage cases (Braak Stages 5-6); PSP – Progressive supranuclear palsy; SD – standard deviation; Galanin- – galanin-negative neurons; Galanin- pTau- – galanin-negative neurons without pTau burden; Galanin- pTau+ – galanin-negative neurons with pTau burden; Galanin+ – galanin neurons; Galanin+ pTau- – galanin neurons without pTau burden; Galanin+ pTau+ – galanin neurons with pTau burden. Comparisons that achieved both statistical significance and sufficient power are highlighted in bold. 1: Wilcoxon rank-sum test; 2: p-values adjusted for multiple comparisons using the Benjamini-Hochberg method; 3: mixed linear regression model accounting for the effects of sex, age at death (AaD), and post-mortem interval (PMI) on the comparisons (e.g., $m0 = \text{measure} \sim \text{group} + \text{Sex} + \text{AaD} + \text{PMI} + (1 | \text{Subject})$ ).

The stereological assessment revealed a reduction in volume and a decrease in the total number of IntN neurons in late-stage PSP and late-AD (*i.e.*, Braak 5 and 6) cases when compared with HC. In PSP cases, we detected an IntN volume reduction of 81.25% (*p* = 0.0115; power = 95.3%) (Figure 3A). Likewise, the neuronal numbers exhibited the most pronounced decline among the analyzed groups, with an 84.91% reduction in neuron count (*p* = 0.0094; power = 100%) (Figure 3B). The PSP neuronal loss is statistically significant and powered even when compared to mid-AD cases (*p* = 0.0241; power = 95.8%) and to late-AD cases (*p* = 0.0095; power = 93.9%) (Table 3). In 3 out of 8 PSP cases (Figure S1), IntN neurons could not be identified despite successfully locating the correct coordinates and expanding the search to all serial sections within the predicted ROI. Thus, the neuronal numbers in these cases were set at zero. Across all PSP cases, a substantial presence of apoptotic debris was observed in the IntN region, supporting the evidence of pronounced neuronal death associated with PSP. All PSP cases were late stage reflecting the relative rarity of PSP and the limited availability of mid- and early-stage cases across brain banks.

**Figure 3.**
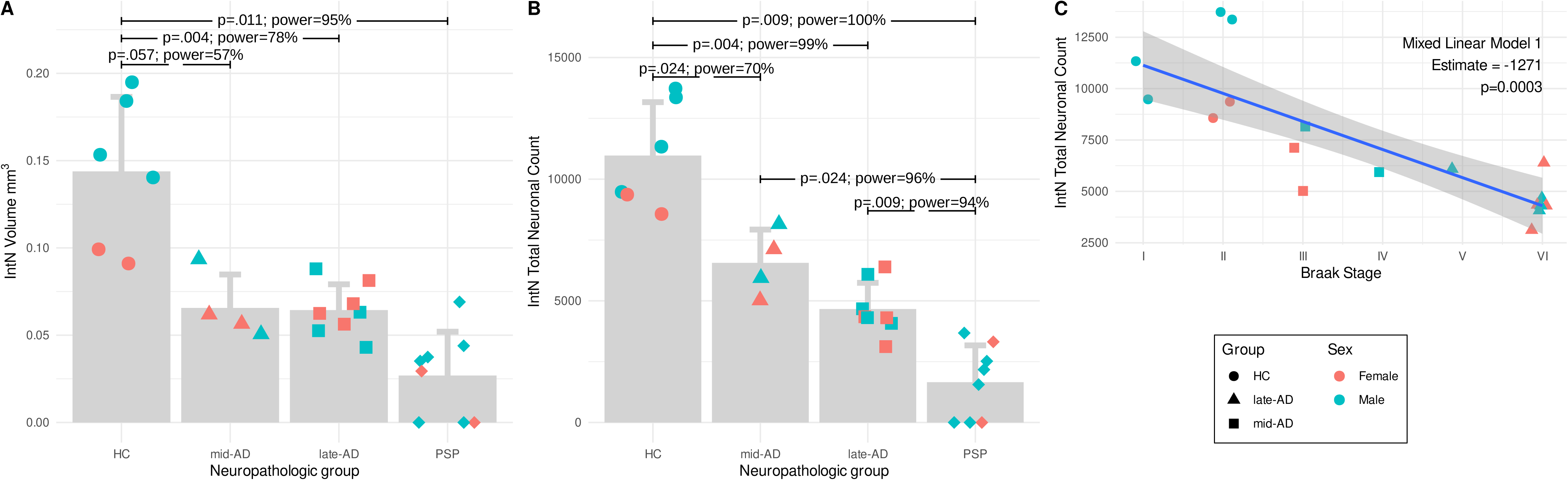
Association Between IntN Volume, Neuronal Count, and Disease Groups or Braak Stage. (A) IntN volume is significantly reduced in mid-AD, late-AD, and PSP compared to HC. Except for mid-AD, all comparisons reached powered statistical significance. (B) Total neuronal count is lower in mid-AD, late-AD, and PSP compared to HC. Additionally, the PSP group shows a significant reduction when compared to both mid- and late-AD. (C) The mixed linear model 1 (see Methods) identifies the Braak stage as the only significant factor associated with neuronal loss in HC and AD subset cases.

### IntN Neurons show a trend to reduction in number already at mid-Braak stages

We observed a 54.38% volume reduction trend (*p* = 0.0571; power = 57.4%; *m*0 *p-value* = 0.0121) when comparing HC with mid-AD cases, but a significant reduction when comparing HC with late-AD (*p* = 0.0040; power = 77.6%). Based on the raw numbers, we hypothesize that volume reduction is likely to occur in mid-AD stages, but the analysis did not reach significance because of the limited sample size (*n* = 4) and lack of power to stratify the analysis by sex, which is desirable because HC male subjects exhibit greater volume compared to females (Figure 3A). The hypothesis of IntN degeneration already at mid-AD stages is supported by an observed 40.21% neuronal reduction in mid-AD stages, which was underpowered (*p* = 0.0241; power = 70.3%), thus requiring confirmation (Figure 3B). Furthermore, the mixed linear model1 (see Methods) identifies the Braak stage as the sole significant factor (*p* < 0.0003) contributing to neuronal reduction in AD cases (Figure 3C).

### Selective neuronal Vulnerability of galaninergic-positive vs negative neurons in IntN

Despite its small size, the IntN has a mixed neuronal population, of which 40% to 50% are galanin-positive. Galanin-positive neurons have been implicated in SWS control in rodents(Kroeger et al., 2018; Szymusiak et al., 1998). Thus, we investigated whether the pattern of neuronal vulnerability of IntN to AD and PSP differed depending on whether the neuron synthetizes galanin. Reflecting the overall neuronal reduction, PSP cases exhibited statistically significant and powered decreases in both subneuronal categories: galanin-positive neurons (- 95.24%; *p* = 0.0115; power = 97.5%) and galanin-negative neurons (−78.63%; *p* = 0.0139; power = 93.6%) compared to HC (Figure 4A-B). Similarly, reductions in galanin-positive and galanin-negative neurons in AD cases paralleled the total neuronal count relative to HC. Galanin-positive neurons exhibited a more pronounced reduction, with decreases of 51.49% in mid-AD (*p* = 0.0381; power = 40.3%) and 76.89% in late-AD (*p* = 0.004; power = 89.4%) (Figure 4A). For galanin-negative neurons, reductions of 33.36% in mid-AD (*p* > 0.05) and 45.72% in late-AD (*p* = 0.0233; power = 52.1%) were observed, with neither achieving sufficient power; however, the latter reached statistical significance (Figure 4B). Slightly lower p-values and higher power were observed using a mixed linear model, though only the late-AD comparison remained statistically significant and powered (Table 3). Despite limitations due to the sample size, the data clearly indicate that galanin-positive neurons are significantly vulnerable to AD-tau, either by dying or losing the ability to synthesize galanin - an aspect this study is not designed to test (Figure 4A).

**Figure 4.**
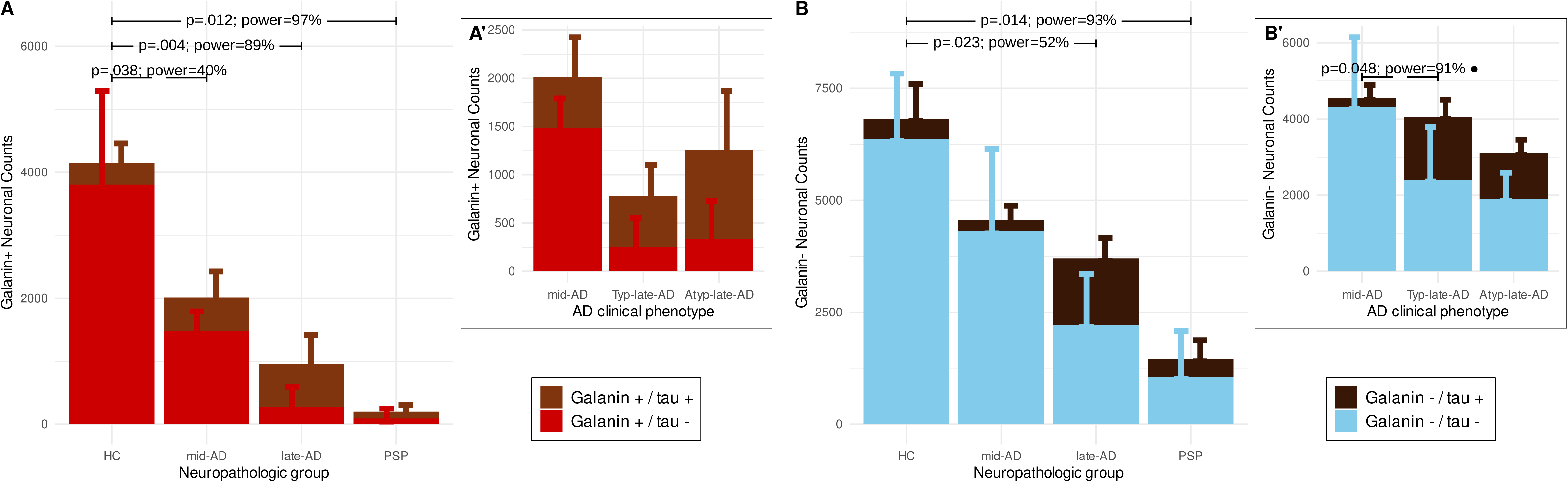
Differential Neuronal Vulnerability of IntN in AD and PSP. (A) Stereological quantification of total galanin-positive neurons across all groups. (B) Stereological quantification of total galanin-negative neurons across all groups. (A’ and B’) highlight differences between typical and atypical AD cases compared to HC. The statistical test presented in B’, close to the ● sign, was applied solely to Galanin- / tau+ neurons and represents the statistical differences in the increase of neuronal counts when the mid-AD cases group is compared with typical AD manifestation cases.

### Phospho-tau burden in galanin-positive vs negative neurons of the IntN

In the Braak stages 0-2 (*i.e.*, HC), p-tau inclusions were sparse, with 9.45% of galanin-positive neurons and 4.82% of galanin-negative neurons showing tau inclusions. By Braak stages 3–4 (*i.e.*, mid-AD), the p-tau burden increased in galanin-positive neurons to 23.49%, while galanin-negative neurons exhibited a burden of 7.29%. In late-AD cases, these numbers reach 76.89% and 42.56% for galanin-positive and -negative neurons, respectively. In PSP, 65.74% of galanin neurons and 29.92% of galanin-negative neurons showed p-tau inclusions. The data suggests an early accumulation of tau aggregates more prominent in galanin neurons than in galanin-negative neurons (Figure 4A-B). In the PSP case, it is hard to evaluate the number of neurons with p-tau due to the massive IntN neuronal loss (Figure 3B).

### Probing differential vulnerability of IntN to AD variants

About 10% of cases meeting neuropathological criteria for AD manifest an atypical, non-amnestic presentation, thus known as atypical AD cases (Graff Radford et al., 2021). A few cases in our late-AD group met the criteria for atypical AD. Given the previous report that atypical AD shows less dysfunction of slow wave sleep than typical AD cases, we ran a relatively unpowered preliminary comparison of typical vs atypical AD, all at late neuropathological stages. IntN volume and total number of neurons were similar in both groups (Table 3). However, two key differences emerged when comparing neuronal subpopulations. First, the number of galanin-positive neurons decreases in typical AD at a higher magnitude than in atypical AD (Figure 4A’). Second, tau burden in galanin-negative neurons increases significantly only in typical AD cases (*p* = 0.048; power = 91%) (Figure 4B’).

## Discussion

The present study provides the first direct quantitative evidence of profound, selective neuronal loss in the human intermediate nucleus (IntN) of the hypothalamus across tauopathies, offering critical insight into disease-specific sleep abnormalities. Early AD-tau accumulation in galanin-positive neurons emerged in cases Braak 0–2 (HC), with a trend toward neuronal loss especially by mid-stage AD (Braak 3–4). This selective vulnerability may explain why sleep disturbances, including impaired SWS, often precede or coincide with clinical AD symptoms. Such early involvement can set off a vicious cycle, as impaired SWS may exacerbate amyloid-β and tau pathology accumulations by reducing sleep-dependent clearance mechanisms (Xie et al., 2013). Notably, we found that PSP, a 4-repeat tauopathy marked by severe sleep initiation and maintenance difficulties (Walsh et al., 2017), exhibits a striking degree of IntN neuronal loss— far exceeding that in late-stage AD. These findings align with our hypothesis that PSP’s inability to maintain stable sleep stems from the near-elimination of SWS-promoting neurons, while wake-promoting circuits remain comparatively intact (J. Oh, Eser, et al., 2019).

Although the literature on the ventrolateral preoptic (VLPO) area in rodents is extensive, research on the human IntN has been limited. Early hypothalamic studies focused primarily on more prominent nuclei, leaving the anterior hypothalamic region less clearly defined. Brockhaus (Brockhaus, 1942) was pivotal in identifying the IntN as a distinct neuronal cluster. Subsequent investigations recognized its putative homology to the VLPO (Allen et al., 1989)(Swaab & Hofman, 1988)(Swaab & Fliers, 1985) but often employed sections stained for Nissl only, complicating precise delineation. Conflicting findings on IntN neuron counts (Allen et al., 1989)(Byne et al., 2000)(LeVay, 1991)(Swaab & Hofman, 1988) highlight the need for reliable chemical markers—such as galanin-immunoreactivity (Gai et al., 1990)—and thicker sections, both incorporated in the present study. Interestingly, although our study was not powered to assess sex differences, we observed a trend toward higher neuron counts in males, similar to previous findings from Swaab and Hofman (Swaab & Hofman, 1988). From a functional neuroanatomy perspective, the IntN’s homology with the VLPO in rodents underscores its probable role in sleep regulation and renewed interest in understanding age and disease-associated changes in the IntN in humans. In rodents, VLPO galaninergic neurons are sleep-active, critical for initiating and maintaining NREM sleep—especially SWS (Kroeger et al., 2018)(Sherin et al., 1996)(Sherin et al., 1998)(Szymusiak et al., 1998). Lesions to the VLPO produce insomnia, while chemogenetic or optogenetic activation of these cells promotes sleep. In humans, Lim et al. (Lim et al., 2014) showed that fewer galaninergic IntN neurons correlate with more fragmented sleep, regardless of the diagnosis of AD. Our findings extend this association into PSP and AD progressive stages, revealing parallel reductions of galanin-positive neurons and p-tau pathology in PSP and AD—both of which frequently present with sleep disturbances.

In AD, progressive tau accumulation in galanin-positive IntN neurons likely compromises their function, leading to early sleep fragmentation and reduced SWS that can precede frank cognitive impairment (Ju et al., 2014). Strikingly, typical AD shows a marked decrease in galanin-positive neurons compared to healthy controls, whereas atypical AD—associated with milder SWS deficits—has relatively preserved galanin-positive cells but exhibits a significant loss of galanin-negative neurons. This divergence supports the notion that galaninergic cell integrity directly influences SWS maintenance, aligning with clinical observations that atypical AD presents fewer disruptions in SWS compared to amnestic AD, although a larger sample size is needed to confirm these preliminary observations.

This study has strengths and limitations. Methodologically, we employed unbiased stereology coupled with immunohistochemistry for galanin and phospho-tau, enabling precise neuron counts and direct assessment of disease-related tau pathology in specific neuronal subtypes. This rigorous approach contrasts with many historical studies that relied on Nissl-stained sections and subjective nuclear boundaries, often confounded by factors such as age, sex, and anatomic orientation (Allen et al., 1989)(Byne et al., 2000)(LeVay, 1991)(Swaab & Hofman, 1988). We selected cases largely free of Lewy bodies or TDP-43 pathology to more confidently attribute changes to the primary tauopathy. However, the rarity of PSP and atypical AD limited sample sizes, restricting generalizability and preventing robust analysis of sex differences. We also lack fine-grained temporal data on precisely when IntN neurons begin to die; future studies examining very early Braak stages (1–2) would clarify whether IntN degeneration predates overt cognitive impairment. Moreover, most cases lacked premortem polysomnography data (J. Y. Oh et al., 2022), limiting direct correlations between neuropathology and clinical sleep phenotypes. Finally, the IntN is not homogeneous: multiple neurochemical phenotypes (Chawla et al., 1997)(Fliers et al., 1994)(Gao & Moore, 1996)(Kruijver et al., 2002) may be distinctly affected, an aspect requiring more granular analysis.

In conclusion, our findings establish that IntN neuronal loss - particularly among galanin-expressing cells - is a key pathological feature in PSP and AD. In PSP, the near-total depletion of these neurons, coupled with relatively intact wake systems, underlies its severe insomnia-like presentation. In AD, the early vulnerability and progressive loss of galanin-positive neurons align with clinical reports of early sleep fragmentation and SWS deficits. Drawing on historical perspectives and addressing methodological controversies, we show that tau pathologies leave distinct “signatures” on these vital sleep-regulating neurons. Further research with larger sample sizes, earlier disease stages, and premortem sleep/wake assessments will refine our understanding and potentially guide therapeutic strategies to preserve restorative sleep in these devastating conditions.

## References

Allen, L. S., Hines, M., Shryne, J. E., & Gorski, R. A. (1989). Two sexually dimorphic cell groups in the human brain. The Journal of Neuroscience, 9(2), 497–506. 10.1523/JNEUROSCI.09-02-00497.1989

Arendt, T., Brückner, M. K., Morawski, M., Jäger, C., & Gertz, H.-J. (2015). Early neurone loss in Alzheimer’s disease: cortical or subcortical? Acta Neuropathologica Communications, 3, 10. 10.1186/s40478-015-0187-1

Attems, J., Toledo, J. B., Walker, L., Gelpi, E., Gentleman, S., Halliday, G., Hortobagyi, T., Jellinger, K., Kovacs, G. G., Lee, E. B., Love, S., McAleese, K. E., Nelson, P. T., Neumann, M., Parkkinen, L., Polvikoski, T., Sikorska, B., Smith, C., Grinberg, L. T., … McKeith, I. G. (2021). Neuropathological consensus criteria for the evaluation of Lewy pathology in post-mortem brains: a multi-centre study. Acta Neuropathologica, 141(2), 159–172. 10.1007/s00401-020-02255-2

Braak, H., & Braak, E. (1991). Neuropathological stageing of Alzheimer-related changes. Acta Neuropathologica, 82(4), 239–259. 10.1007/BF00308809

Brockhaus, H. (1942). Beitrag zur normalen anatomie des hypothalamus und der zona incerta beim menschen. J Psychol Neurol, 51, 96–196.

Byne, W., Lasco, M. S., Kemether, E., Shinwari, A., Edgar, M. A., Morgello, S., Jones, L. B., & Tobet, S. (2000). The interstitial nuclei of the human anterior hypothalamus: an investigation of sexual variation in volume and cell size, number and density. Brain Research, 856(1–2), 254–258. 10.1016/s0006-8993(99)02458-0

Chawla, M. K., Gutierrez, G. M., Young, W. S., McMullen, N. T., & Rance, N. E. (1997). Localization of neurons expressing substance P and neurokinin B gene transcripts in the human hypothalamus and basal forebrain. The Journal of Comparative Neurology, 384(3), 429–442. 10.1002/(sici)1096-9861(19970804)384:3<429::aid-cne8>3.0.co;2-5

Ehrenberg, A. J., Kelberman, M. A., Liu, K. Y., Dahl, M. J., Weinshenker, D., Falgàs, N., Dutt, S., Mather, M., Ludwig, M., Betts, M. J., Winer, J. R., Teipel, S., Weigand, A. J., Eschenko, O., Hämmerer, D., Leiman, M., Counts, S. E., Shine, J. M., Robertson, I. H., … Grinberg, L. T. (2023). Priorities for research on neuromodulatory subcortical systems in Alzheimer’s disease: Position paper from the NSS PIA of ISTAART. Alzheimer’s & Dementia, 19(5), 2182–2196. 10.1002/alz.12937

Falgàs, N., Walsh, C. M., Yack, L., Simon, A. J., Allen, I. E., Kramer, J. H., Rosen, H. J., Joie, R. L., Rabinovici, G., Miller, B., Spina, S., Seeley, W. W., Ranasinghe, K., Vossel, K., Neylan, T. C., & Grinberg, L. T. (2023). Alzheimer’s disease phenotypes show different sleep architecture. Alzheimer’s & Dementia, 19(8), 3272–3282. 10.1002/alz.12963

Fliers, E., Noppen, N. W., Wiersinga, W. M., Visser, T. J., & Swaab, D. F. (1994). Distribution of thyrotropin-releasing hormone (TRH)-containing cells and fibers in the human hypothalamus. The Journal of Comparative Neurology, 350(2), 311–323. 10.1002/cne.903500213

Gai, W. P., Geffen, L. B., & Blessing, W. W. (1990). Galanin immunoreactive neurons in the human hypothalamus: colocalization with vasopressin-containing neurons. The Journal of Comparative Neurology, 298(3), 265–280. 10.1002/cne.902980302

Gao, B., & Moore, R. Y. (1996). The sexually dimorphic nucleus of the hypothalamus contains GABA neurons in rat and man. Brain Research, 742(1–2), 163–171. 10.1016/s0006-8993(96)01005-0

Garcia-Falgueras, A., Ligtenberg, L., Kruijver, F. P. M., & Swaab, D. F. (2011). Galanin neurons in the intermediate nucleus (InM) of the human hypothalamus in relation to sex, age, and gender identity. The Journal of Comparative Neurology, 519(15), 3061–3084. 10.1002/cne.22666

Graff-Radford, J., Yong, K. X. X., Apostolova, L. G., Bouwman, F. H., Carrillo, M., Dickerson, B. C., Rabinovici, G. D., Schott, J. M., Jones, D. T., & Murray, M. E. (2021). New insights into atypical Alzheimer’s disease in the era of biomarkers. Lancet Neurology, 20(3), 222–234. 10.1016/S1474-4422(20)30440-3

Grinberg, L. T., Ferretti, R. E. de L., Farfel, J. M., Leite, R., Pasqualucci, C. A., Rosemberg, S., Nitrini, R., Saldiva, P. H. N., Filho, W. J., & Brazilian Aging Brain Study Group. (2007). Brain bank of the Brazilian aging brain study group - a milestone reached and more than 1,600 collected brains. Cell and Tissue Banking, 8(2), 151–162. 10.1007/s10561-006-9022-z

Heinsen, H., Arzberger, T., & Schmitz, C. (2000). Celloidin mounting (embedding without infiltration) - a new, simple and reliable method for producing serial sections of high thickness through complete human brains and its application to stereological and immunohistochemical investigations. Journal of Chemical Neuroanatomy, 20(1), 49–59.

Höglinger, G. U., Respondek, G., Stamelou, M., Kurz, C., Josephs, K. A., Lang, A. E., Mollenhauer, B., Müller, U., Nilsson, C., Whitwell, J. L., Arzberger, T., Englund, E., Gelpi, E., Giese, A., Irwin, D. J., Meissner, W. G., Pantelyat, A., Rajput, A., van Swieten, J. C., … Movement Disorder Society-endorsed PSP Study Group. (2017). Clinical diagnosis of progressive supranuclear palsy: The movement disorder society criteria. Movement Disorders, 32(6), 853–864. 10.1002/mds.26987

Horner, R. L., & Peever, J. H. (2017). Brain circuitry controlling sleep and wakefulness. Continuum (Minneapolis, Minn.), 23(4, Sleep Neurology), 955–972. 10.1212/CON.0000000000000495

Iranzo, A. (2016). Sleep in neurodegenerative diseases. Sleep Medicine Clinics, 11(1), 1–18. 10.1016/j.jsmc.2015.10.011

Ju, Y.-E. S., Lucey, B. P., & Holtzman, D. M. (2014). Sleep and Alzheimer disease pathology--a bidirectional relationship. Nature Reviews. Neurology, 10(2), 115–119. 10.1038/nrneurol.2013.269

Kaalund, S. S., Passamonti, L., Allinson, K. S. J., Murley, A. G., Robbins, T. W., Spillantini, M. G., & Rowe, J. B. (2020). Locus coeruleus pathology in progressive supranuclear palsy, and its relation to disease severity. Acta Neuropathologica Communications, 8(1), 11. 10.1186/s40478-020-0886-0

Kroeger, D., Absi, G., Gagliardi, C., Bandaru, S. S., Madara, J. C., Ferrari, L. L., Arrigoni, E., Münzberg, H., Scammell, T. E., Saper, C. B., & Vetrivelan, R. (2018). Galanin neurons in the ventrolateral preoptic area promote sleep and heat loss in mice. Nature Communications, 9(1), 4129. 10.1038/s41467-018-06590-7

Kruijver, F. P. M., Balesar, R., Espila, A. M., Unmehopa, U. A., & Swaab, D. F. (2002). Estrogen receptor-alpha distribution in the human hypothalamus in relation to sex and endocrine status. The Journal of Comparative Neurology, 454(2), 115–139. 10.1002/cne.10416

LeVay, S. (1991). A difference in hypothalamic structure between heterosexual and homosexual men. Science, 253(5023), 1034–1037. 10.1126/science.1887219

Lim, A. S. P., Ellison, B. A., Wang, J. L., Yu, L., Schneider, J. A., Buchman, A. S., Bennett, D. A., & Saper, C. B. (2014). Sleep is related to neuron numbers in the ventrolateral preoptic/intermediate nucleus in older adults with and without Alzheimer’s disease. Brain: A Journal of Neurology, 137(Pt 10), 2847–2861. 10.1093/brain/awu222

Moran, M., Lynch, C. A., Walsh, C., Coen, R., Coakley, D., & Lawlor, B. A. (2005). Sleep disturbance in mild to moderate Alzheimer’s disease. Sleep Medicine, 6(4), 347–352. 10.1016/j.sleep.2004.12.005

Oh, J., Eser, R. A., Ehrenberg, A. J., Morales, D., Petersen, C., Kudlacek, J., Dunlop, S. R., Theofilas, P., Resende, E. D. P. F., Cosme, C., Alho, E. J. L., Spina, S., Walsh, C. M., Miller, B. L., Seeley, W. W., Bittencourt, J. C., Neylan, T. C., Heinsen, H., & Grinberg, L. T. (2019). Profound degeneration of wake-promoting neurons in Alzheimer’s disease. Alzheimer’s & Dementia, 15(10), 1253–1263. 10.1016/j.jalz.2019.06.3916

Oh, J., Petersen, C., Walsh, C. M., Bittencourt, J. C., Neylan, T. C., & Grinberg, L. T. (2019). The role of co-neurotransmitters in sleep and wake regulation. Molecular Psychiatry, 24(9), 1284–1295. 10.1038/s41380-018-0291-2

Oh, J. Y., Walsh, C. M., Ranasinghe, K., Mladinov, M., Pereira, F. L., Petersen, C., Falgàs, N., Yack, L., Lamore, T., Nasar, R., Lew, C., Li, S., Metzler, T., Coppola, Q., Pandher, N., Le, M., Heuer, H. W., Heinsen, H., Spina, S., … Grinberg, L. T. (2022). Subcortical neuronal correlates of sleep in neurodegenerative diseases. JAMA Neurology, 79(5), 498–508. 10.1001/jamaneurol.2022.0429

Pina-Escudero, S. D., La Joie, R., Spina, S., Hwang, J.-H., Miller, Z. A., Huang, E. J., Grant, H., Mundada, N. S., Boxer, A. L., Gorno-Tempini, M. L., Rosen, H. J., Kramer, J. H., Miller, B. L., Seeley, W. W., Rabinovici, G. D., & Grinberg, L. T. (2024). Comorbid neuropathology and atypical presentation of Alzheimer’s disease. *Alzheimer’s & Dementia*□*: Diagnosis*, Assessment & Disease Monitoring, 16(3), e12602. 10.1002/dad2.12602

Polsinelli, A. J., & Apostolova, L. G. (2022). Atypical alzheimer disease variants. *Continuum (Minneapolis*, Minn.), 28(3), 676–701. 10.1212/CON.0000000000001082

Saper, C. B., & Fuller, P. M. (2017). Wake-sleep circuitry: an overview. Current Opinion in Neurobiology, 44, 186–192. 10.1016/j.conb.2017.03.021

Saper, C. B. (2021). The intermediate nucleus in humans: Cytoarchitecture, chemoarchitecture, and relation to sleep, sex, and Alzheimer disease. Handbook of Clinical Neurology, 179, 461–469. 10.1016/B978-0-12-819975-6.00030-3

Satpati, A., Pereira, F. L., Soloviev, A., Mladinov, M., Leite, R. E. P., Suemoto, C. K., Rodriguez, R. D., Paes, V. R., Walsh, C. M., Spina, S., Seeley, W. W., Pasquallucci, C. A., Jacob□Filho, W., Neylan, T. C., & Grinberg, L. T. (2023). Deciphering the molecular signature of selective neuronal vulnerability in the wake□promoting lateral hypothalamic area in Alzheimer’s disease: A digital multiplexed gene expression study across Braak stages. Alzheimer’s & Dementia, 19(S12). 10.1002/alz.079466

Sherin, J. E., Elmquist, J. K., Torrealba, F., & Saper, C. B. (1998). Innervation of histaminergic tuberomammillary neurons by GABAergic and galaninergic neurons in the ventrolateral preoptic nucleus of the rat. The Journal of Neuroscience, 18(12), 4705–4721. 10.1523/JNEUROSCI.18-12-04705.1998

Sherin, J. E., Shiromani, P. J., McCarley, R. W., & Saper, C. B. (1996). Activation of ventrolateral preoptic neurons during sleep. Science, 271(5246), 216–219. 10.1126/science.271.5246.216

Slomianka, L., & West, M. J. (2005). Estimators of the precision of stereological estimates: an example based on the CA1 pyramidal cell layer of rats. Neuroscience, 136(3), 757–767. 10.1016/j.neuroscience.2005.06.086

Spina, S., La Joie, R., Petersen, C., Nolan, A. L., Cuevas, D., Cosme, C., Hepker, M., Hwang, J.-H., Miller, Z. A., Huang, E. J., Karydas, A. M., Grant, H., Boxer, A. L., Gorno-Tempini, M. L., Rosen, H. J., Kramer, J. H., Miller, B. L., Seeley, W. W., Rabinovici, G. D., & Grinberg, L. T. (2021). Comorbid neuropathological diagnoses in early versus late-onset Alzheimer’s disease. Brain: A Journal of Neurology, 144(7), 2186–2198. 10.1093/brain/awab099

Suemoto, C. K., Ferretti-Rebustini, R. E. L., Rodriguez, R. D., Leite, R. E. P., Soterio, L., Brucki, S. M. D., Spera, R. R., Cippiciani, T. M., Farfel, J. M., Chiavegatto Filho, A., Naslavsky, M. S., Zatz, M., Pasqualucci, C. A., Jacob-Filho, W., Nitrini, R., & Grinberg, L. T. (2017). Neuropathological diagnoses and clinical correlates in older adults in Brazil: A cross-sectional study. PLoS Medicine, 14(3), e1002267. 10.1371/journal.pmed.1002267

Swaab, D. F., & Fliers, E. (1985). A sexually dimorphic nucleus in the human brain. Science, 228(4703), 1112– 1115. 10.1126/science.3992248

Swaab, D. F., & Hofman, M. A. (1988). Sexual differentiation of the human hypothalamus: ontogeny of the sexually dimorphic nucleus of the preoptic area. Brain Research. Developmental Brain Research, 44(2), 314–318.

Szymusiak, R., Alam, N., Steininger, T. L., & McGinty, D. (1998). Sleep-waking discharge patterns of ventrolateral preoptic/anterior hypothalamic neurons in rats. Brain Research, 803(1–2), 178–188. 10.1016/s0006-8993(98)00631-3

Thal, D. R., Rüb, U., Orantes, M., & Braak, H. (2002). Phases of A beta-deposition in the human brain and its relevance for the development of AD. Neurology, 58(12), 1791–1800. 10.1212/wnl.58.12.1791

Theofilas, P., Ehrenberg, A. J., Nguy, A., Thackrey, J. M., Dunlop, S., Mejia, M. B., Alho, A. T., Paraizo Leite, R. E., Rodriguez, R. D., Suemoto, C. K., Nascimento, C. F., Chin, M., Medina-Cleghorn, D., Cuervo, A. M., Arkin, M., Seeley, W. W., Miller, B. L., Nitrini, R., Pasqualucci, C. A., … Grinberg, L. T. (2018). Probing the correlation of neuronal loss, neurofibrillary tangles, and cell death markers across the Alzheimer’s disease Braak stages: a quantitative study in humans. Neurobiology of Aging, 61, 1–12. 10.1016/j.neurobiolaging.2017.09.007

Theofilas, P., Polichiso, L., Wang, X., Lima, L. C., Alho, A. T. L., Leite, R. E. P., Suemoto, C. K., Pasqualucci, C. A., Jacob-Filho, W., Heinsen, H., Brazilian Aging Brain Study Group, & Grinberg, L. T. (2014). A novel approach for integrative studies on neurodegenerative diseases in human brains. Journal of Neuroscience Methods, 226, 171–183. 10.1016/j.jneumeth.2014.01.030

Tsuneoka, Y., & Funato, H. (2021). Cellular composition of the preoptic area regulating sleep, parental, and sexual behavior. Frontiers in Neuroscience, 15, 649159. 10.3389/fnins.2021.649159

Walsh, C. M., Ruoff, L., Walker, K., Emery, A., Varbel, J., Karageorgiou, E., Luong, P. N., Mance, I., Heuer, H. W., Boxer, A. L., Grinberg, L. T., Kramer, J. H., Miller, B. L., & Neylan, T. C. (2017). Sleepless night and day, the plight of progressive supranuclear palsy. Sleep, 40(11). 10.1093/sleep/zsx154

Wang, S., Zheng, X., Huang, J., Liu, J., Li, C., & Shang, H. (2024). Sleep characteristics and risk of Alzheimer’s disease: a systematic review and meta-analysis of longitudinal studies. Journal of Neurology, 271(7), 3782–3793. 10.1007/s00415-024-12380-7

West, M. J. (2013). Tissue shrinkage and stereological studies. Cold Spring Harbor Protocols, 2013(3). 10.1101/pdb.top071860

Xie, L., Kang, H., Xu, Q., Chen, M. J., Liao, Y., Thiyagarajan, M., O’Donnell, J., Christensen, D. J., Nicholson, C., Iliff, J. J., Takano, T., Deane, R., & Nedergaard, M. (2013). Sleep drives metabolite clearance from the adult brain. Science, 342(6156), 373–377. 10.1126/science.1241224

