## Supplementary Material for "Tau-Associated Neuronal Loss in the Intermediate Nucleus of the Human Hypothalamus (VLPO Analog): Unveiling the Basis of NREM Sleep Dysfunction in PSP and Alzheimer’s Disease"

**Table S1. Individual demographics data and unbiased stereology counts.**

| Subject | Sex | Age at Death | Disease Duration | PMI | Brain Weight (g) | Clinical Diagnosis | Neuropat-hology Diagnosis | Braak Stage | CERAD | | Thal | | CDR | | CDR SOB | | ABC Score | | IntN volume (mm^3^) | IntN neuronal numbers | CE counts |
| --- | --- | --- | --- | --- | --- | --- | --- | --- | --- | --- | --- | --- | --- | --- | --- | --- | --- | --- | --- | --- | --- |
| 1 | M | 53 | - | 21.23 | 1548 | None | HC | 1 | 0 | | 0 | | 0 | | 0 | | A0B1C0 | | 0.184171 | 11334.73 | 0.03 |
| 2 | M | 66 | - | 10.55 | 1283 | None | HC | 1 | 0 | | 0 | | 0 | | 0 | | A0B1C0 | | 0.140374 | 9477.03 | 0.03 |
| 3 | M | 58 | - | 11.48 | 1402 | None | HC | 2 | 0 | | 0 | | 0 | | NA | | A0B1C0 | | 0.153377 | 13361.26 | 0.02 |
| 4 | F | 67.9 | - | 15.5 | 1.196 | None | HC | 2 | 0 | | 0 | | 0.5 | | 1.5 | | A0B1C0 | | 0.091085 | 8567.73 | 0.03 |
| 5 | F | 83 | - | 12.13 | 1020 | None | HC | 2 | 0 | | 1 | | 0 | | 0 | | A1B1C0 | | 0.0992138 | 9364.62 | 0.04 |
| 6 | M | 73 | - | 18.75 | 1488 | None | HC | 2 | 0 | | 1 | | 0 | | 0 | | A1B1C0 | | 0.194934 | 13725.83 | 0.02 |
| 7 | M | 73 | NA | 17 | 1140 | AD | AD | 3 | 0 | | 0 | | 1 | | 8 | | A0B2C0 | | 0.0505911 | 8155.78 | 0.03 |
| 8 | F | 67 | NA | 14.4 | 1332 | None | AD | 3 | 0 | | 0 | | 0 | | 0 | | A0B2C0 | | 0.0618587 | 7120.93 | 0.04 |
| 9 | F | 68 | NA | 13.6 | 1206 | None | AD | 3 | 0 | | 0 | | 0 | | 0 | | A0B2C0 | | 0.056531 | 5021.29 | 0.04 |
| 10 | M | 81 | 7 | 28.5 | 1375 | MCI | AD | 4 | 0 | | 0 | | NA | | NA | | A0B2C0 | | 0.093523 | 5939.67 | 0.04 |
| 11 | M | 78 | 15 | NTD | 1060 | CBS | AD | 6 | 3 | | 5 | | 3 | | NA | | A3B3C3 | | 0.0525735 | 4072.45 | 0.04 |
| 12 | M | 58 | 10 | 7.47 | 1231 | lvPPA | AD | 6 | 3 | | 5 | | 3 | | 18 | | A3B3C3 | | 0.0631608 | 4674.25 | 0.05 |
| 13 | F | 62 | 6 | 15.33 | 1230 | PCA | AD | 6 | 3 | | 5 | | 3 | | 17 | | A3B3C3 | | 0.0813477 | 4344.59 | 0.04 |
| 14 | M | 94 | NA | 12.17 | 936 | AD | AD | 5 | NA | | NA | | 1 | | 5 | | NA | | 0.0880408 | 6090.63 | 0.05 |
| 15 | F | 81 | NA | 21.1 | 1160 | AD | AD | 6 | 2 | | 2 | | 2 | | 12 | | A1B3C2 | | 0.0563 | 6392.62 | 0.04 |
| 16 | F | 56 | 8 | 4.92 | 986 | AD | AD | 6 | 3 | | 5 | | 3 | | NA | | A3B3C3 | | 0.062516 | 4303.96 | 0.04 |
| 17 | F | 60 | 7 | 13 | 1011 | AD | AD | 6 | 3 | | 5 | | 3 | | 17 | | A3B3C3 | | 0.0680627 | 3119.29 | 0.05 |
| 18 | M | 64 | 5 | 23.67 | 1335 | AD | AD | 6 | 3 | | 5 | | 3 | | 18 | | A3B3C3 | | 0.0430566 | 4306.74 | 0.04 |
| 19 | M | 63 | 13 | 4.42 | 1365 | PSP-S | PSP | 1 | 0 | | 2 | | 3 | | 17 | | A1B1C0 | | 0 | 0 | - |
| 20 | F | 66 | 9 | 24.78 | 930 | PSP-S | PSP | 2 | 0 | | 0 | | 0.5 | | 11 | | A0B1C0 | | 0 | 0 | - |
| 21 | M | 59 | 7 | 6 | 1292 | PSP-S | PSP | 1 | 0 | | 2 | | 1 | | 8 | | A1B1C0 | | 0.0439348 | 2520.06 | 0.06 |
| 22 | M | 68 | 6 | 10.08 | 1250 | PSP-S | PSP | 0 | 0 | | 0 | | 0.5 | | 11 | | A0B0C0 | | 0.0374843 | 1557.7 | 0.07 |
| 23 | M | 75 | 13 | NA | 1239 | PSP-S | PSP | 2 | 0 | | 1 | | 3 | | 13 | | A1B1C0 | | 0.0690414 | 3676.43 | 0.04 |
| 24 | M | 61 | 3 | NA | 1547 | PSP-S | PSP | 2 | 1 | | 2 | | 2 | | 14 | | A1B1C1 | | 0 | 0 | - |
| 25 | M | 76 | 10 | 8.25 | 1204 | PSP-S | PSP | 3 | 2 | | 5 | | 0.5 | | 6 | | A3B2C2 | | 0.0351692 | 2172.3 | 0.06 |
| 26 | F | 81 | 9 | 15.28 | 951 | nfvPPA | PSP | 3 | 2 | 5 | | 3 | | 18 | | A3B2C2 | | 0.0294344 | | 3314.66 | 0.06 |

Abbreviations: PMI – postmortem interval; g – grams; CDR: Clinical Dementia Rating; SOB – sums of boxes; mm3 - cubic millimeter; CE counts - Coefficient of Error for neuronal number counts; M – Male; F – Female; NA – not accessed; HC – Healthy control; AD – Alzheimer’s disease; MCI - Mild cognitive impairment; CBS – Corticobasal Syndrome; lvPPA – Logopenic variant primary progressive aphasia; PCA – Posterior cortical atrophy; PSP: Progressive supranuclear palsy; PSP-S – PSP syndrome; nfvPPA - Nonfluent variant primary progressive aphasia.

**Table S2. Demographics stratified by clinical phenotypical manifestation of AD.**

| Characteristics | HC | Typ-late-AD | Atyp-late-AD | p-value |
| --- | --- | --- | --- | --- |
| N | 6 | 5 | 3 |  |
| Age at Death, mean (SD), y | 66.8 (10.68) | 71 (16) | 66 (10.58) | 0.652^t^ |
| Males, No. (%) | 4 (66.7) | 2 (40) | 2 (66.7) | 1^C^ |
| Disease duration, mean (SD) | 0 (0) | 6.7 (1.53) | 10.3 (4.51) | 0.275^K^ |
| Braak Stage, range | 1-2 | 5-6 | 6-6 | 0.0339^C^ |
| CDR-SOB, median (IQR) | 0 (0) | 14.5 (7) | 17.5 (0.5) | 0.475^W^ |
| Brain weight, mean (SD), g | 1123.7 (580.75) | 1085.6 (162.48) | 1173.7 (98.44) | 0.393^W^ |
| PMI, mean (SD), h | 14.9 (4.33) | 15 (7.52) | 11.4 (5.56) | 0.393^W^ |

Abbreviations: HC – Healthy controls; Typ-late-AD – Amnestic Alzheimer’s disease cases (Braak Stages 5-6); Atyp-late-AD – Atypical Alzheimer’s disease cases (Braak Stages 5-6); N – number of individuals; SD – standard deviation; IQR – interquartile range; PMI – postmortem interval. P-values were calculated conforming data type, normality, and homogeneity of variance: t – t-Test; C – chi-square; K – Kruskal-Wallis; W – Wilcoxon rank-test.

**Table S3. Mean (SD) and percentage of stereological estimates for different characteristics of the IntN stratified by clinical phenotypical manifestation of AD.**

| Characteristics | HC | Typ-late-AD | Atyp-late-AD |
| --- | --- | --- | --- |
| Volume, mean (SD), mm3 | 0.144 (0.0427) | 0.064 (0.0165) | 0.066 (0.0146) |
| Neuron population, mean (SD), un | 10971.9 (2191.93) | 4842.7 (1369.96) | 4363.8 (301.36) |
| Sub-neuronal population (n) |  |  |  |
| Galanin-, mean (SD) | 6824.8 (2157.17) | 4062.6 (1358.58) | 3108.2 (976.9) |
| Galanin- pTau-, mean (SD) | 6373.8 (1454.69) | 2406.7 (1376.71) | 1893.9 (695.36) |
| Galanin- pTau+, mean (SD) | 451 (775.7) | 1655.9 (448.19) | 1214.4 (350.03) |
| Galanin+, mean (SD) | 4147.1 (1412.93) | 780.1 (579.98) | 1255.5 (1003.19) |
| Galanin+ pTau-, mean (SD) | 3803.9 (1480.95) | 252.6 (302.66) | 331.6 (399.44) |
| Galanin+ pTau+, mean (SD) | 343.2 (310.58) | 527.4 (321.92) | 923.9 (616.15) |
| Sub-neuronal population (%) |  |  |  |
| Percentage of Galanin-, mean (SD) | 61.82 (11.6) | 83.23 (11.4) | 71.43 (23.1) |
| Percentage of Galanin- pTau-, mean (SD) | 95.18 (7.1) | 55.77 (16.6) | 60.22 (5.8) |
| Percentage of Galanin- pTau+, mean (SD) | 4.82 (7.1) | 44.23 (16.6) | 39.78 (5.8) |
| Percentage of Galanin+, mean (SD) | 38.18 (11.6) | 16.77 (11.4) | 28.57 (23.1) |
| Percentage of Galanin+ pTau-, mean (SD) | 90.55 (10.3) | 24.39 (22.2) | 20.99 (10.8) |
| Percentage of Galanin+ pTau+, mean (SD) | 9.45 (10.3) | 75.61 (22.2) | 79.01 (10.8) |

Abbreviations: HC – Healthy controls; AD – Alzheimer’s disease; Typ-late-AD – Amnestic Alzheimer’s disease cases (Braak Stages 5-6); Atyp-late-AD – Atypical Alzheimer’s disease cases (Braak Stages 5-6); SD – standard deviation; Galanin- – galanin-negative neurons; Galanin- pTau- – galanin-negative neurons without pTau burden; Galanin- pTau+ – galanin-negative neurons with pTau burden; Galanin+ – galanin neurons; Galanin+ pTau- – galanin neurons without pTau burden; Galanin+ pTau+ – galanin neurons with pTau burden.

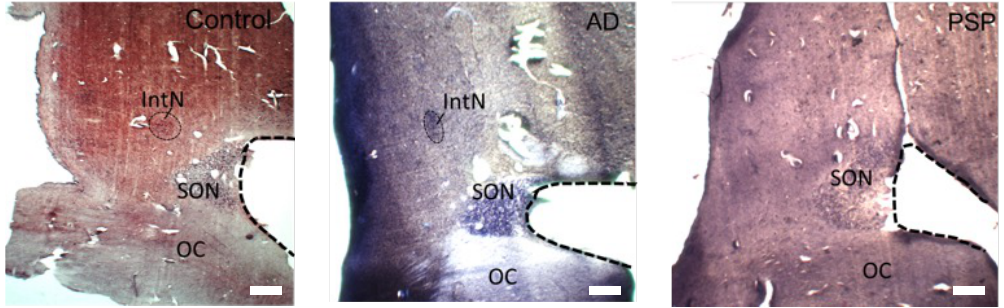

**Figure S1. A histological section containing the IntN ROI*.*** The IntN is located near the crossing of the anterior commissure and lateral to the supraoptic and suprachiasmatic nuclei. The identification was done using both gross anatomical landmarks and histological cues. Scale (white bar on the bottom right) bars: 1 mm. The third column, PSP case, depicts the disappearance of the ROI at naked eye examination in PSP. Supraoptic nucleus (SON) and Optic chiasm (OC) are demonstrated as the regional landmarks of the preoptic area.Sciwheel inserting bibliography...
